# Role of NLRP3 activation in salt sensitive blood pressure regulation, effect of ND-13

**DOI:** 10.1101/2025.02.26.640392

**Authors:** Raisha Garcia, Jacob Polzin, Celia Arias, Estela Guillén, Patricia Lathan, María Del Carmen Gallego-López, Carmen De Miguel, Jun Feranil, Hewang Lee, Ines Armando, Pedro Jose, Santiago Cuevas

## Abstract

**Background and Objectives:** High salt intake is a major contributor to the development and exacerbation of hypertension, partly by inducing an inflammatory response through immune cell dysfunction. Inflammasomes, key components of the innate immune response, may influence blood pressure regulation. The renal DJ-1 protein is known for its antioxidant and anti-inflammatory properties. To explore novel pharmacological applications of renal DJ-1 pathway, we developed ND-13, a peptide consisting of 13 highly conserved amino acids derived from the DJ-1 sequence. In this study, we investigated the effects of ND-13 and MCC950, a specific NLRP3 inflammasome inhibitor, on blood pressure regulation in C57BL/6J mice on a high-salt diet.

**Methods:** C57BL/6J mice were fed a high-salt diet (HS) for one week and then treated with ND-13 or MCC950, an NLRP3 inflammasome inhibitor. Subsequently, gene expression by qPCR, staining of immune cells, Sirius Red and Periodic Acid-Schiff (PAS) staining were determined in the mice kidneys, as well as the inflammasome activity in peritoneal cells.

**Results:** One week of HS resulted increased in blood pressure, that was prevented by both ND-13 and MCC950 treatments. These treatments also prevented the increase in proteinuria that was accompanied by tubular protein deposits. Renal expression of inflammatory genes, immune cell infiltration, and renal collagen deposition were not observed in the HS group. Peritoneal macrophages isolated from HS treated mice exhibited enhanced IL-1β release upon LPS+ATP stimulation, suggesting activation of the NLRP3 inflammasome. Treatment with ND-13 and MCC950 normalized this activity. Furthermore, ND-13 reduced IL-1β mRNA expression in peritoneal macrophages.

**Conclusions:** Our findings highlight the critical role of the NLRP3 inflammasome in salt-sensitive blood pressure regulation and suggest that ND-13 may serve as a potential therapeutic agent for preventing hypertension and associated inflammatory alterations induced by a high salt intake.

## 1. Introduction

Salt sensitivity refers to a condition where an individual’s blood pressure (BP) significantly increases in response to high salt (NaCl) intake. Salt sensitivity is linked to genetics, age, obesity, kidney function, and other factors. It plays a critical role in the risk of developing hypertension and cardiovascular diseases (1). Inflammation is a key contributor to salt sensitivity, as it impairs sodium regulation and elevates BP. Chronic inflammation affects renal function by reducing the kidneys’ ability to excrete excess sodium, while also promoting vascular stiffness and dysfunction. These effects amplify the BP response to high salt intake, underscoring the critical role of inflammation in the pathogenesis and progression of salt-sensitive hypertension (2, 3).

The NLRP3 inflammasome, a protein complex within the innate immune system, detects danger signals and triggers inflammatory responses by activating pro-inflammatory cytokines like IL-1β and IL-18. Its activation has been implicated in various inflammatory and cardiovascular diseases (4). Regarding BP regulation, studies have demonstrated that pharmacological inhibition of the NLRP3 inflammasome reduces BP, renal damage, and dysfunction in salt-sensitive hypertension (5). Moreover, activation of the NLRP3 inflammasome in vascular smooth muscle cells induces phenotypic changes that impair vascular function and contribute to the increase in BP (6).

The DJ-1 protein serves as an oxidative stress sensor with diverse functions, including transcriptional regulation and antioxidant activities. In the kidneys, DJ-1 plays a protective role by mitigating oxidative stress and inflammation (6), and it is involved in regulating podocyte apoptosis and mitophagy through the DJ-1/PTEN pathway, maintaining the normal morphology, structure, and function of glomerular podocytes (7). Previous studies from our group have demonstrated that renal DJ-1 exhibits antioxidant and anti-inflammatory properties (6, 8). Mice with kidney-specific silencing of DJ-1 or with germline deletion of DJ-1 (DJ-1-/-) exhibit elevated BP and reduced expression and activity of Nrf2, a master regulator of antioxidant and anti-inflammatory responses (8). These findings suggest that DJ-1 inhibits renal reactive oxygen species (ROS) production, at least in part, via activation of the Nrf2-antioxidant gene pathway. Additionally, activation of the DJ-1/Nrf2 pathway has been implicated in the pathogenesis of diabetic nephropathy in rats (9). Collectively, these observations highlight the essential role of DJ-1 in preserving renal function and protecting against kidney injury.

A peptide based on the amino acid sequence of the DJ-1 protein was developed by Dr. Offen’s team. They identified the most conserved region of the protein, consisting of a 13-amino-acid sequence, and fused it with a 7-amino-acid cell-penetrating peptide to ensure cellular uptake. The resulting 20-amino-acid compound was named ND-13. Multiple studies have shown that ND-13 protects neuronal cultures from Parkinson’s disease related neurotoxins and other pathological conditions (10-12). ND-13 reduces cell death and deactivates the pro-apoptotic enzyme caspase-3 in neuronal cells subjected to neurotoxic damage. Additionally, ND-13 activates the Nrf2 pathway, leading to the upregulation of antioxidant genes regulated by Nrf2, such as heme oxygenase-1, NAD(P)H quinone dehydrogenase 1 (NQO1), Glutamate-cysteine ligase <u>catalytic subunit</u> (GCLC), and Glutamate-cysteine ligase modifier subunit (GCLM) (11, 13) These findings are consistent with our previous observations in the kidney (8).

This study aimed to determine the potential protective effects of ND-13 and the DJ-1/Nrf2 pathway on the blood pressure response to high salt intake.

## 2. Material and Methods

### Animals

In this project, we used a C57Bl/6J (Jackson Laboratories, Bar Harbor, ME) murine model to investigate the effects of NLRP3 inflammasome activation and inhibition on renal inflammation and BP. We and others have already reported that the C57Bl/6J strain is salt sensitive (14). Male mice (8 weeks old) were randomly assigned to four groups, all mice were fed normal salt diet (NS, 0.8% NaCl) then three groups were shifted to high salt diet (HS, 4% NaCl) for 7 days which coincided with the duration of treatment with saline solution, ND-13 (3 mg/kg/day), and MCC950 (10 mg/kg/day) given by intraperitoneal injection. Urine (24-hr output) was collected while mice where on NS and again on day 7 of treatment while on HS using MMC100 metabolic cages for mice (Hatteras Instruments, Grantsboro, NC). All animal manipulations were approved by the George Washington University Institutional Animal Care and Use Committee (A2022-014).

### Measurement of Blood Pressure by Tail-Cu8 Plethysmography

The CODA Monitor noninvasive blood pressure acquisition system for mice (Kent Scientific, Torrington, CT) was used for all tail-cu^ measurements. This method uses a specialized volume pressure recording (VPR) sensor, to detect blood pressure based on volume changes in the tail. Blood pressure (BP) was measured while mice were fed NS diet before shifting to HS diet (serving as control BP) and at the end of the treatment period on HS diet. Briefly, mice were anesthetized with isoflurane (1-3%; 28-37 ml/min flow rate based on body weight and pre-determined by the system) using SomnoSuite low-flow anesthesia delivery system designed specifically for mice (Kent Scientific, Torrington, CT) and placed on the SurgiSuite multi-functional surgical platform with integrated far infrared warming pad (Kent Scientific, Torrington, CT). Respiration was visually monitored. The occlusion cu^ was placed at the base of the tail and the VPR sensor cuff was placed adjacent to the occlusion cuff. The temperature at the base of the tail is kept between 32-35 degree C monitored using a handheld infrared thermometer. After a 5-minute equilibration period, pressure readings were acquired for 10–13 consecutive minutes. The VPR sensor cu^ detects changes in the tail volume as the blood returns to the tail during the occlusion cu^ deflation. Each recording session consisted of 15 to 25 inflation and deflation cycles per set, of which the first 5 cycles were “acclimation” cycles and were not used in the analysis, whereas the following cycles were used. At the end of the experiment, mice were sacrificed and biospecimens were collected for cellular and molecular studies. Leukocytes were collected by intraperitoneal irrigation and aspiration.

### Methods

Several assays were performed to evaluate the inflammatory responses and renal function. We determined secreted IL-1β and tumor necrosis factor-alpha (TNF-α) in both leukocyte medium and serum, LDH assays to measure cytotoxicity or pyroptosis rates, urinary microalbumin, and creatinine assays to assess kidney function through the microalbumin-to-creatinine ratio, and qRT-PCR was performed for several transcripts related to cytokines, inflammasome regulation, and fibrosis in renal cortices and leukocyte lysates.

IL1β (Invitrogen, 88-7013A) and TNF-α (88-7324, Invitrogen) and urime microalbumin (80630, Crystal Chem) were measured by ELISA kits following the protocols described by the manufacturers. Urinary creatinine was measured by a colorimetric assay (500701, Cayman Chemical).

### qPCR Assays

RNA was extracted from mouse renal cortex and leukocyte lysates using the RNeasy Kit (Qiagen, 74104) according to the manufacturer’s instructions. RNA was quantified by spectrophotometry and the volume adjusted accordingly. cDNA was synthesized using the PrimeScript RT Reagent Kit with gDNA Eraser (RR047A, Takara) and quantitative PCR was performed on an iQ 5 Real-Time PCR System (BioRad) using SYBR Green Mastermix (Thermo Fisher) according to the manufacturer’s guidelines. Prevalidated RT2 qPCR primers (Qiagen) were used to amplify mouse genes of interest, including Tnf-α (NM_013693. 3), Il-6 (NM_001314054), Il-18 (NM_008360), Il-1β (NM_008361), Tgfβ1 (NM_011577), Nlrp3 (NM_145827), Pycard (NM_023258), Casd1 (NM_145398), P2rx7 (NM_011027) and Col4a5 (NM_007736). Gapdh (NM_008084) was used as a control transcript and data were analyzed using the ΔΔCt method.

### Preparation of Paraffin-Embedded Renal Tissue

Dissected kidney tissue samples were fixed in 10% neutral buffered formalin (CAS: 50-00-0, HT501128, MiliporeSigma) for 48 hours, embedded in paraffin and sectioned using a microtome (Leica RM2125 RTS, Leica Biosystems) to obtain 4-10 μm thick sections.

### Histologic Methods

Kidney pathology was evaluated in H&E stained slides in a blinded fashion with comparisons among four groups. Slide sections were used for histologic analyses after whole slide scanning (Aperio CS2) and viewed on ImageScope (Leica Aperio). Renal pathology was assessed for evidence of inflammatory infiltrates and injury to vasculature, glomeruli and tubules. Five cortical glomeruli per mouse were randomly selected from the mid-cortex on the basis of a roughly round contour and the number of cells and area of the capillary tuft portion of the glomerulus (not including Bowmans space) was measured using the annotation tool available in SlideViewer (3-D Histech). The cell counts in the glomeruli are reported as an average for the mice in each group. The areas of the five largest hilar glomeruli per mouse were also evaluated with measurement of the capillary tuft glomerular area as well as the glomerular area including Bowman’s space. The significance of measurements between groups was determined by one-way ANOVA with Tukey post-hoc testing

### Histological Analysis of Fibrosis Quantification

Renal fibrosis was assessed using Sirius Red staining, and images were acquired using bright-field microscopy (Olympus BX40 equipped with a 10X eyepiece lens; Olympus America, Melville, NY). Whole-kidney scans were generated using a motorized XY stage and a digital camera (Olympus DP71). Sequential 20X magnification images were captured and digitally reconstructed using CellSense imaging software (Olympus). The cortical region of each kidney scan was delineated, and the percentage of fibrotic deposition (stained blue) within the outlined area was quantified using MetaMorph software (Molecular Devices LLC., San Jose, CA). The mean percentage of fibrotic area for each experimental group was calculated and normalized to the control group.

### Immunohistochemical Analysis and Quantification of Immune Cell Infiltration

Renal tissue sections were incubated with primary antibodies targeting CD3 (1:600 dilution; Abcam, Cambridge, MA) and F4/80 (1:200 dilution; Bio-Rad, Hercules, CA), followed by detection with a polymer-conjugated secondary antibody (Biocare Medical, Concord, CA). T-lymphocyte infiltration (CD3+ cells) was quantified by blindly counting 10 microscopic fields (200 x 200 μm, 400X magnification) within the renal cortex of each sample. The average number of infiltrating T cells per field was calculated and reported. Macrophage infiltration (F4/80+ cells) was assessed by capturing 10 cortical images per animal at 400X magnification. The percentage of cortical area positive for F4/80 staining was quantified using MetaMorph software, and the mean expression per animal was determined. Data are presented as the average percentage of F4/80-positive area per experimental group.

### Periodic Acid-Schiff (PAS) Staining of Tissue Slides

Tissue slides were subjected to Periodic Acid-Schiff (PAS) staining to assess carbohydrate-rich structures such as glycogen, glycoproteins, and basement membranes. Slides were immersed in Periodic Acid Solution (CAS: 10450-60-9, 3951, Sigma-Aldrich) rinsed thoroughly and incubated in Schiff’s Reagent (CAS: 569-61-9, 3952016, Sigma-Aldrich). Hematoxylin was used to counterstain nuclei. The slides were scanned using a Leica SCN400 (Leica Microsystems), the images were analyzed using the Leica Microsystem Viewer (Leica Microsystems) and the percentage of the staining was quantified for each sample and compared with the control.

### Leukocytes culture and treatments

The peritoneal cell suspension in PBS was centrifuged at 300xg for eight minutes and the supernatant was aspirated. The cell pellet was then washed with a further 5 mL of PBS (10010049, Gibco), spun again and the supernatant discarded. The cell pellet was resuspended in 1 mL of RPMI medium (1640, Sigma-Aldrich), the viable cells in the suspension were counted by haemocytometry and 0.32% trypan blue (15250061, Gibco) and transferred to 24-well cell culture plates. The cells were then incubated for 2-3 hours at 37°C in a 5% CO2 incubator to induce adherence to the plate. The wells were aspirated, washed once with PBS and incubated for 24 hours in RPMI medium supplemented with 10% fetal bovine serum (FBS) (A5670701, Gibco), 1% L-glutamine (A2916801, Gibco) and 1% penicillin-streptomycin (10,000 U/mL) (15140122, Gibco).

The collected leukocytes were first incubated with various treatments: control, 2 μg/mL lipopolysaccharide (LPS), adenosine triphosphate (ATP) 3 mmol/L, or LPS+ATP to prime and induce inflammasome activity, respectively. The incubation protocol included the addition of LPS of a control approximately 24 hours after plating, followed by the addition of ATP or a control three hours later. After 20 minutes of incubation with ATP or control, medium was collected from all wells and cryopreserved, RNA was extracted from adherent leukocytes The LDH cytotoxicity assay (11644793001, Roche) was used to determine the concentration of lactate dehydrogenase (LDH) released from damaged leukocytes into the media relative to the LDH remaining in intact leukocytes collected from adherent cells using a Triton X-100 detergent solution. The percent cytotoxicity was calculated to estimate the pyroptotic potential of each sample by subtracting the absorbance of the respective control-treated media from that of the LPS- or LPS+ATP-treated cell media and lysates.

### Statistical Analysis

Data is presented as means ± SE. Comparisons between two groups were performed using Student’s t-test. For comparisons among three or more groups one-way ANOVA was performed using Prism9 (GraphPad Software, Boston, MA), followed by Fisher’s least significant difference (LSD), Holm-Sidak test or Tuckey tests to determine significant pairwise differences. A p-value of less than 0.05 was considered statistically significant, where * (*p < 0.033*), ** (*p < 0.002*), and *** (*p < 0.001*).

## 3. Results

### 3.1 Effect of ND-13 and MCC950 treatment on blood pressure, systemic inflammation and renal function in C57Bl/6J mice fed a high salt diet

The increased in the salt content of the diet increased systolic BP only in mice treated with saline. Both ND-13 and MCC950 treatments resulted in a significant reduction in BP compared to both , their BPs on NS and that of mice on HS+saline, demonstrating their effectiveness in mitigating the BP effects of a HS (**Fig. 1A)**. Unexpectedly, MCC950 treatment led to a slight, yet significant, elevation in serum IL-1β levels, contrary to its known function as an NLRP3 inhibitor. This outcome may suggest the presence of a compensatory feedback mechanism that maintains baseline IL-1β levels despite inflammasome inhibition (**Fig. 1B**). The urine microalbumin-to-creatinine ratio increased nonsignificantly in all treatment groups, except for the ND-13 group, following the implementation of the HS diet but remained within a healthy range. It appears that neither ND-13 nor MCC950 could considerably reduce this rise, indicating that under the experimental conditions, their BP-lowering benefits do not extend to reducing the urinary microalbumin-to-creatinine ratio (**Fig. 1C**).

**Figure 1.**
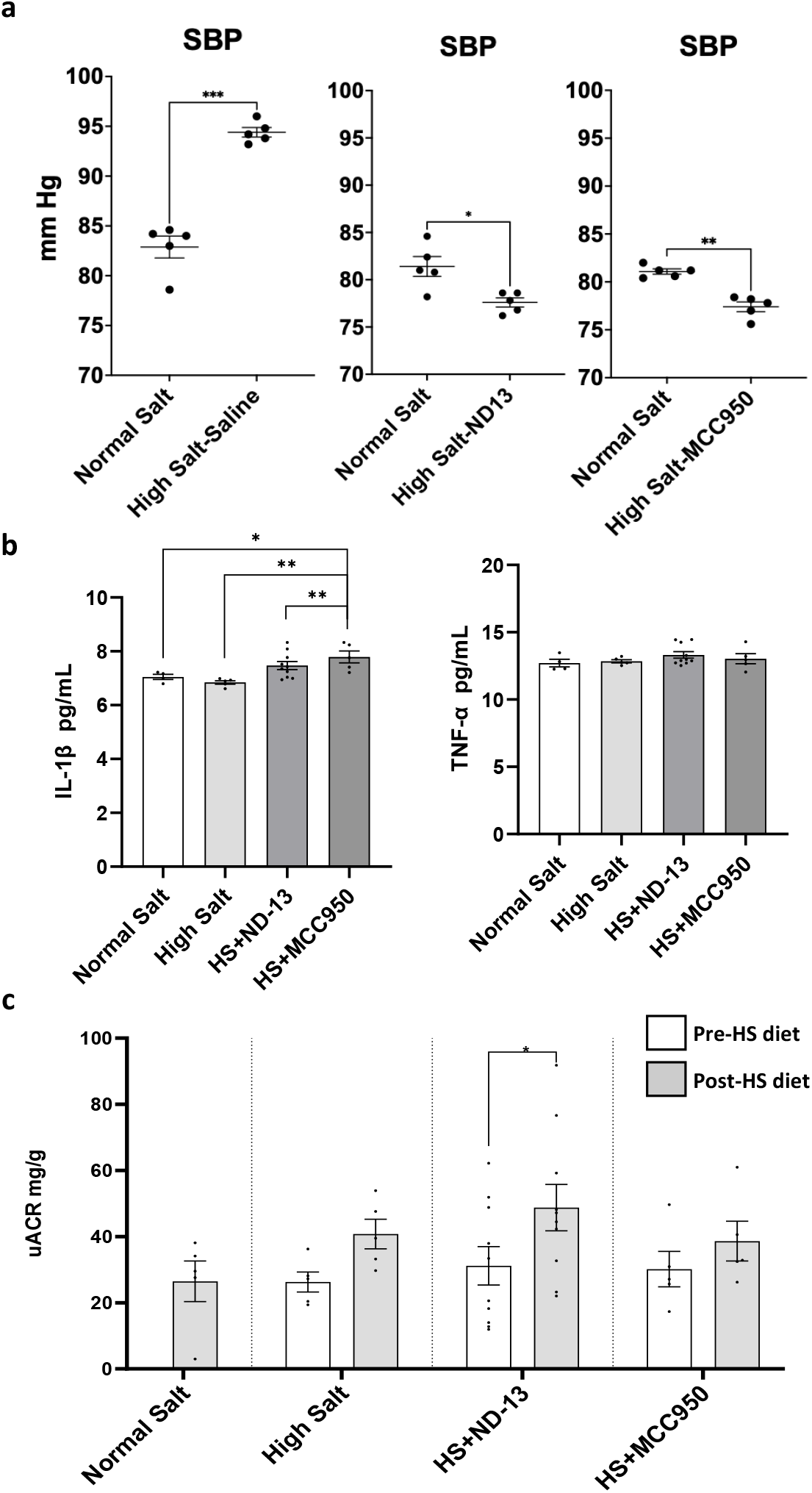
Systolic blood pressure levels, IL-1β and TNF-α plasma levels and proteinuria in mice with high salt diet. **(a)** Effect of ND-13 on systolic blood pressure in mice high salt diet-induced. **(b)** Release of IL-1β and TNF-α in serum of mice high salt diet-induced (NS n=4, HS n=5, HS+ND-13 n=10, HS+MCC950 n= 5). **(c)** Urinary microalbumin-to-creatinine ratio of mice high salt diet-induced (n=5). Data is represented as mean ± SEM; one-way ANOVA was used in b and c; significance levels are indicated as follow: \**p<0.05;* \*\**p<0.01*.

### 3.2 Effect of ND-13 and MCC950 treatment on fibrosis and immune cells renal infiltration in C57Bl/6J mice fed HS

Analysis of Sirius Red staining showed no effect of HS diet on renal fibrosis. However, ND-13 had a decreasing trend, consistent with our previous publication demonstrating the effect of ND-13 on renal damage (15)(**Fig. 2a**). In both cases, the population of T-lymphocytes (CD3+ cells) and macrophages (F4/80+ cells) in the cortex showed a tendency to decrease in response to a high-salt diet. However, this interpretation remains speculative, as no significant differences were found between groups **(Fig. 2b-c**). MCC950 treatment significantly increased the number of macrophages (F4/80+ cells), possibly due to the inhibition of pyroptosis, a cell death mechanism induced by inflammasome activation. (**Fig. 2c**).

**Figure 2.**
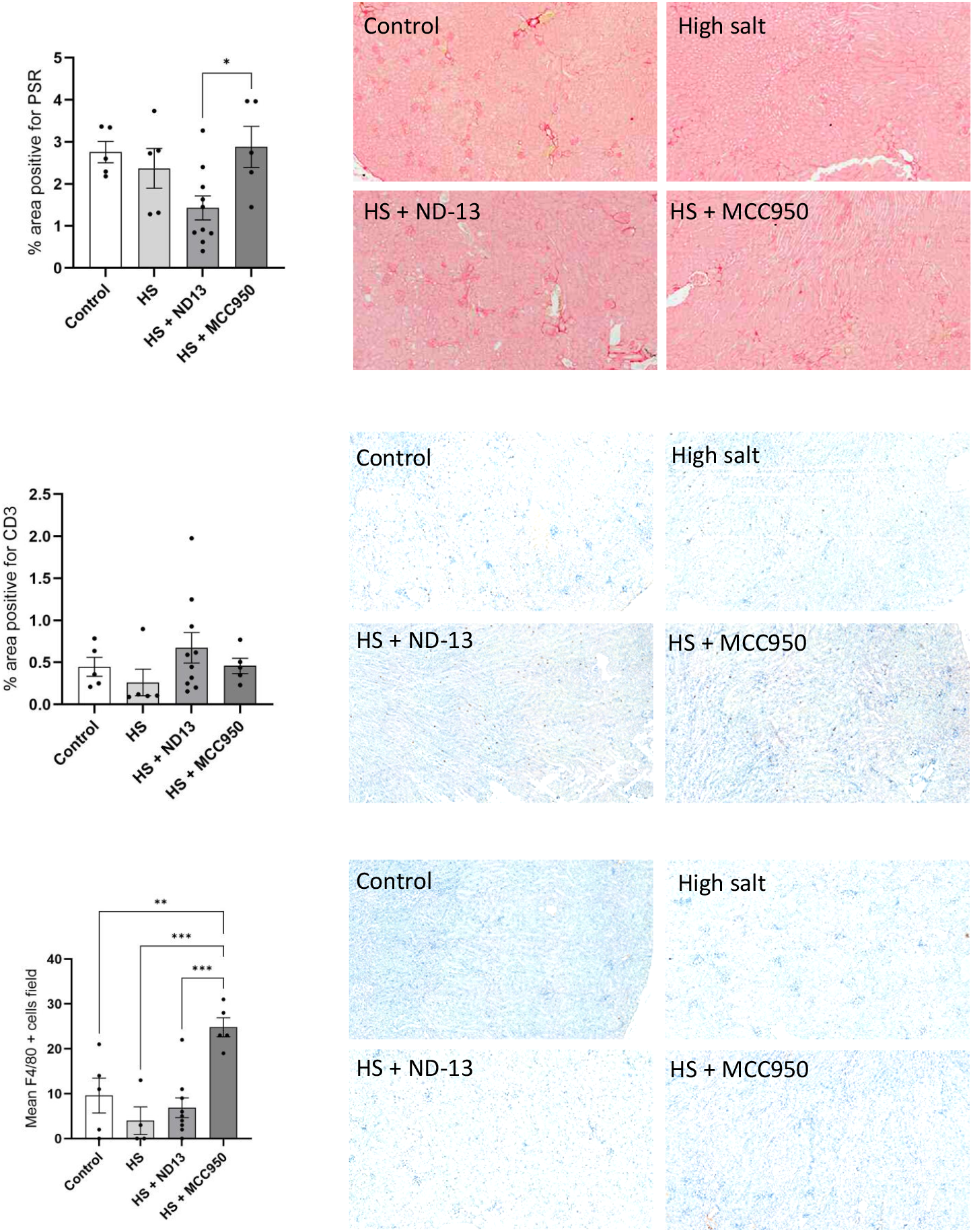
Sirius Red straining quantification data and macrophages and T cell infiltration, into the kidney cortex in mice on high salt diet. A) Quantification of collagen deposition in the renal cortex of C57BL/6 mice subjected to a high-salt diet and treated with ND-13 and MCC950. (B-C) Representative images and quantification of T-cell (CD3+ cells) and macrophage (F4/80+ cells) infiltration in the renal cortex of C57BL/6 mice under a high-salt diet with ND-13 and MCC950 treatment. *n* = 5 per group; *p < 0.05* vs. sham, two-way Tukey’s test.

### 3.3 Effect of ND-13 and MCC950 treatment on glomerular and tubular damage determined by PAS staining quantification in C57Bl/6J mice fed HS

Quantification of PAS staining in glomeruli and renal tubules of mice on a high-salt diet revealed a clear tendency toward increased deposition, particularly in tubules of the renal cortex. Glomerular damage quantification of PAS staining was also elevated. Treatment with ND-13 and MCC950 effectively prevented the increased accumulation of carbohydrate macromolecules, such as glycogen, and mucosubstances, including glycoproteins, glycolipids, and mucins, in renal tissues. This reduction is associated with the prevention of renal damage (**Fig. 3**). No significant pathology was seen in any of the groups of mice on routine sections of kidneys stained with H&E. There was a tendency for greater volume in Bowman’s space within hilar glomeruli, but the difference was not statistically significant between the groups of mice in the mid-cortical glomeruli or when comparing measurements of area in the five largest glomeruli in each group. (**Table 1**).

**Figure 3.**
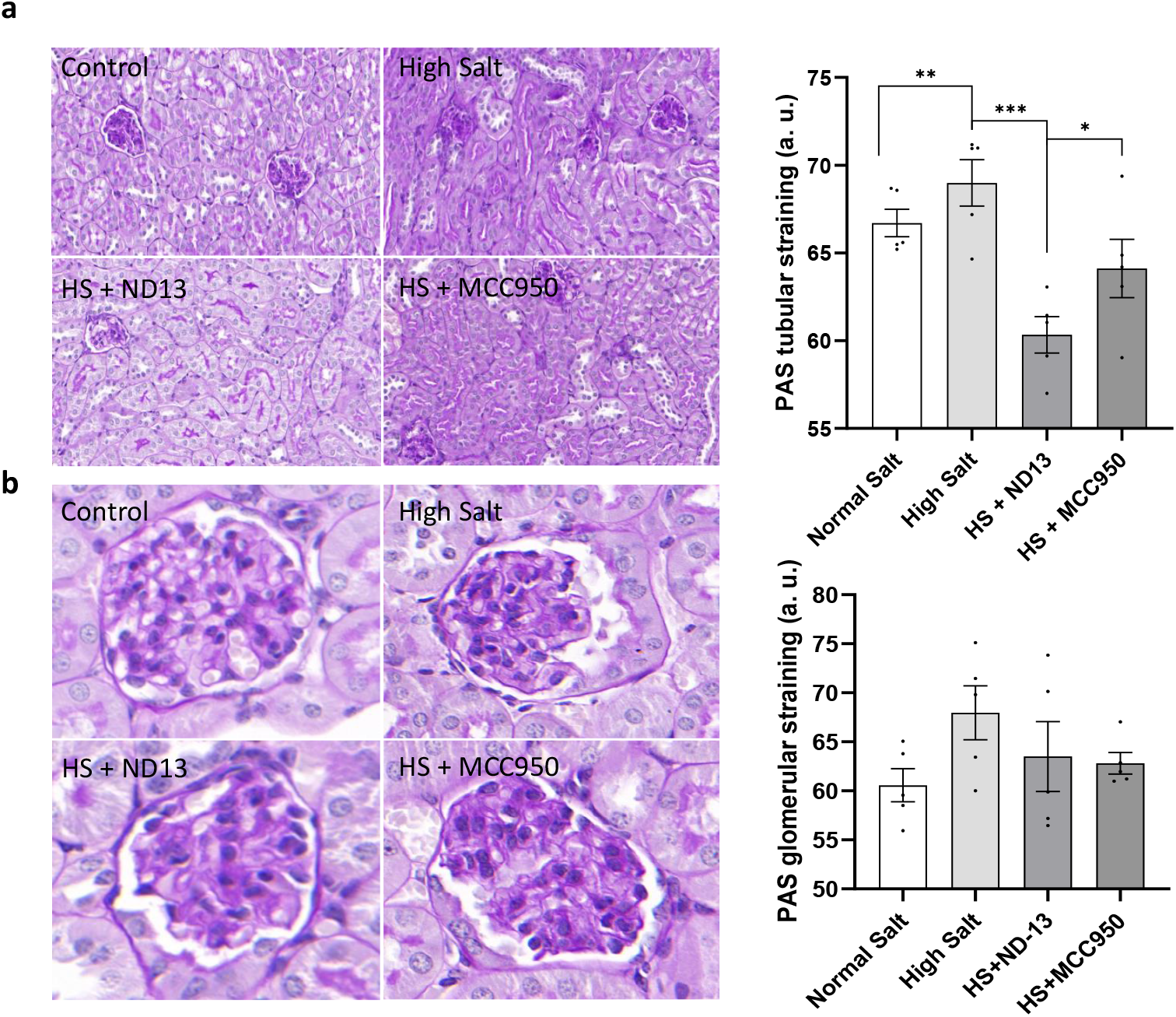
PAS straining quantification data in glomeruli and renal tubules of mice on high salt diet. **(a)** Periodic Acid Schiff (PAS) staining to detect tubular protein deposits and assess tubular structural abnormalities: five right and left random kidney sections per mouse, respectively, in 40X fields, with several tubules in each, n=5 animals per group. **(b)** Periodic Acid Schiff (PAS) staining to detect glomerular protein deposits and assess glomerular structural abnormalities: five right and left random kidney sections per mouse, respectively, in 63X fields, n=5 animals per group. Data is represented as mean ± SEM; one-way ANOVA multiple comparisons was used in a and b; significance levels are indicated as follow: *\*p<0*.*05*; *\*\*p<0*.*005*; *\*\*\*p<0*.*0005*; no differences were found in b (*p>0*.*05*).

**Table 1.**
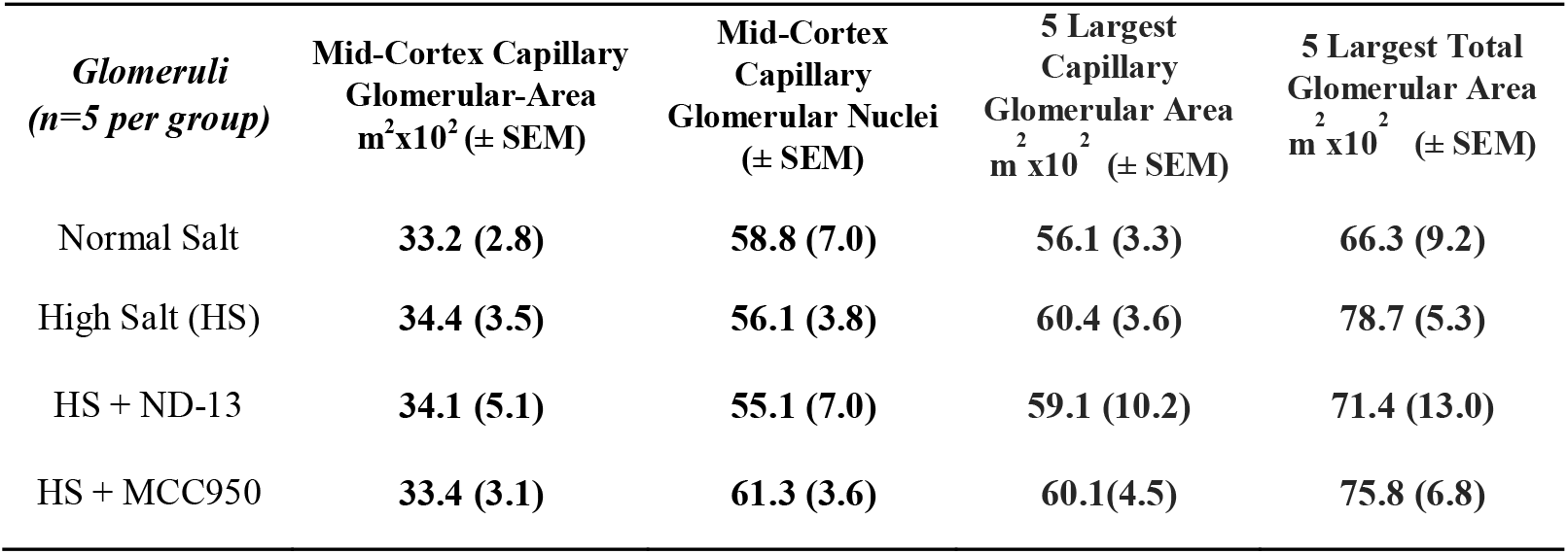
Area of Mid-Cortex Capillary Glomeruli and the Five Largest Capillary Glomeruli in Mice on a Normal Salt Diet vs. a High-Salt Diet With or Without Treatment. There are no statistically significant differences between group means as determined by one-way ANOVA for Capillary Glomerular Area (F 3,21)= 0.0763, p = 0.9721 and Glomerular nuclei (F 3,21) = 0.8649, p = 0.4747. HSD Tukey post-hoc testing found no significant differences between any of the groups. There are no statistically significant differences between group means as determined by one-way ANOVA for Capillary Total Glomerular area (F 3,21)= 0.2230, p = 0.8793 and Glomerular area including Bowman’s capsule (F 3,21)= 0.9389, p = 0.4395. HSD Tukey post-hoc testing found no significant differences between any of the groups.

| *Glomeruli*<br>(*n=5 per group*) | Mid-Cortex Capillary<br>Glomerular-Area<br>$\text{m}^2 \times 10^2 (\pm \text{SEM})$ | Mid-Cortex<br>Capillary<br>Glomerular Nuclei<br>($\pm \text{SEM}$) | 5 Largest<br>Capillary<br>Glomerular Area<br>$\text{m}^2 \times 10^2 (\pm \text{SEM})$ | 5 Largest Total<br>Glomerular Area<br>$\text{m}^2 \times 10^2 (\pm \text{SEM})$ |
| --- | --- | --- | --- | --- |
| Normal Salt | 33.2 (2.8) | 58.8 (7.0) | 56.1 (3.3) | 66.3 (9.2) |
| High Salt (HS) | 34.4 (3.5) | 56.1 (3.8) | 60.4 (3.6) | 78.7 (5.3) |
| HS + ND-13 | 34.1 (5.1) | 55.1 (7.0) | 59.1 (10.2) | 71.4 (13.0) |
| HS + MCC950 | 33.4 (3.1) | 61.3 (3.6) | 60.1(4.5) | 75.8 (6.8) |

### 3.4 Effect of ND-13 and MCC950 treatment on mRNA renal cortex expression of target genes in C57BL/6J mice fed a high-salt diet

qRT-PCR of renal cortices was performed to analyze transcripts related to cytokines, inflammasome regulation, and renal fibrosis. The results from the renal cortex revealed that *Tgfβ1*, a key gene associated with fibrosis, and *Col4a5* showed no significant differences in transcription among the groups.

Interestingly, the renal transcription of *Il-6* and *Tnf-α* was significantly reduced in mice fed HS compared to mice on NS. No significant differences in *Il-6* transcription were detected between untreated HS diet mice and those treated with MCC950. However, normalization of *Il-6* and *Tnf-α* transcription to NS levels was observed in ND-13-treated mice compared to untreated high-salt diet mice (**Fig.4b**). This normalization may result from compensatory mechanisms induced by homeostasis dysregulation. Previous studies have described the potential antihypertensive effects of *IL-6* and *TNF-α* expression in the renal cortex during high-salt diets (16), suggesting that inflammation associated with increased blood pressure under HS dietary conditions is independent of NFκB signaling.

**Figure 4.**
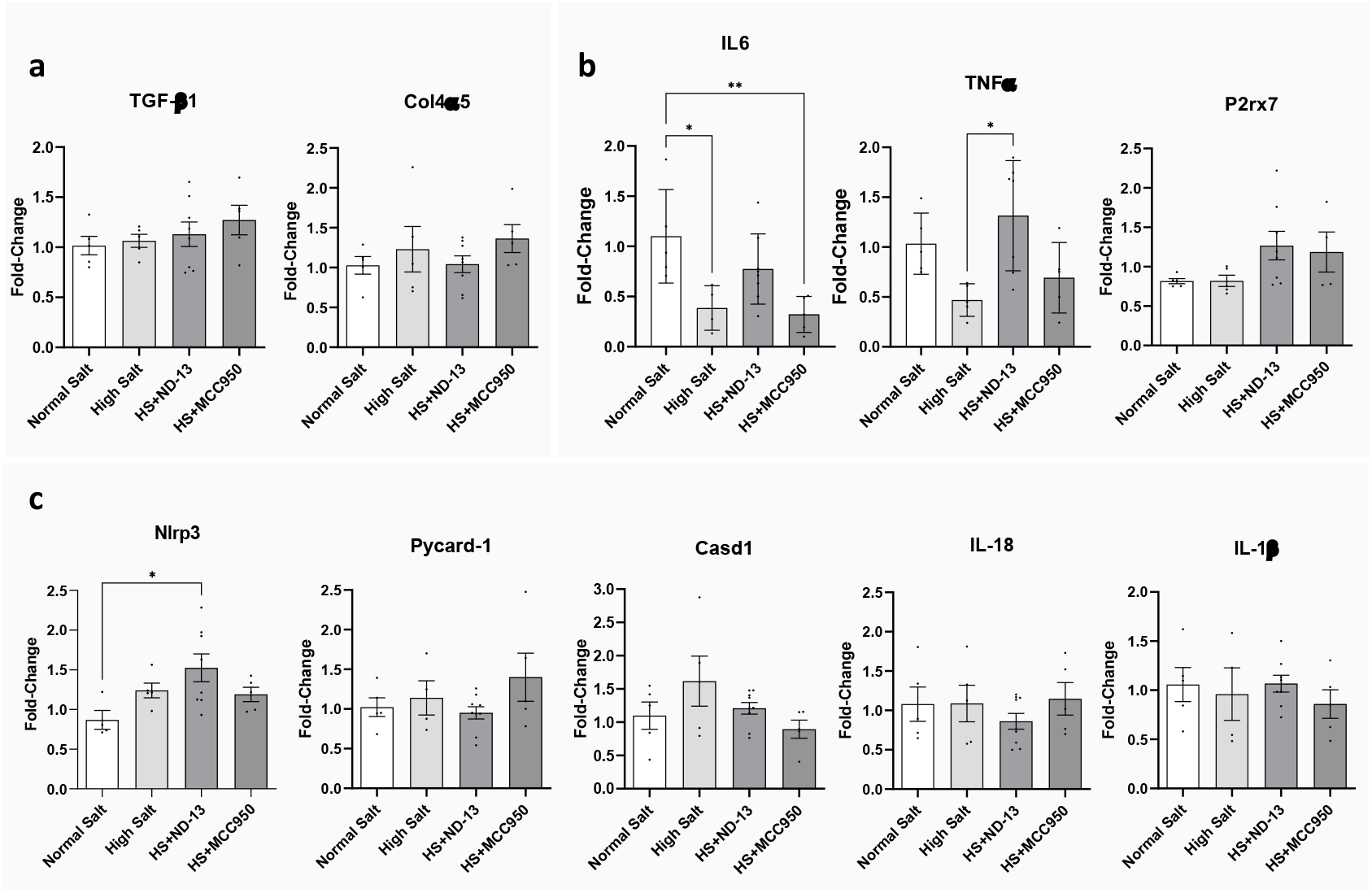
mRNA expression in the renal cortex. **(a)** mRNA expression in the renal cortex of genes associated with fibrosis. **(b)** mRNA expression in the renal cortex of genes associated with inflammatory. **(c)** mRNA expression in the renal cortex of genes associated with inflammasome regulation. Data is represented as mean ± SEM; one-way ANOVA was used in a-c; significance levels are indicated as follow: \**p<0.05*, \*\**p<0.01* (NS n=5, HS n=5, HS+ND-13 n=9, HS+MCC950 n=5).

In contrast, the transcription of genes involved in inflammasome regulation, including *P2rx7, Nlrp3, Pycard-1, Casp1, Il-18*, and *Il-1*β, did not vary significantly between control mice on a normal-salt diet, mice on a high-salt diet, and mice treated with either ND-13 or MCC950. However treatment with ND-13 did increase *Nlrp3* expression suggesting an enhanced inflammatory response. (**Fig.4c**).

### 3.5 Effect of ND-13 and MCC950 treatment on NLRP3 inflammasome activation and mRNA expression of target genes in peritoneal immune cells of mice fed HS

Peritoneal immune cells were cultured in the laboratory and activated with LPS and ATP to assess inflammasome NLRP3 activation. IL-1β secretion requires two-step stimulation: priming with a pathogen-associated molecular pattern (PAMP), such as <u>LPS</u>, and triggering with a damage-associated molecular pattern (DAMP), in this case, <u>ATP</u> (see **Fig. 5**). Supernatant analysis revealed significant findings regarding IL-1β and TNF-α secretion by leukocytes. IL-1β release is specifically associated with inflammasome activation, in contrast to TNF-α levels, which were elevated in LPS-treated groups due to LPS-induced TNF-α production via NFkB activation (for pathways involved in NLRP3 activation, see Figure 5)

**Figure 5.**
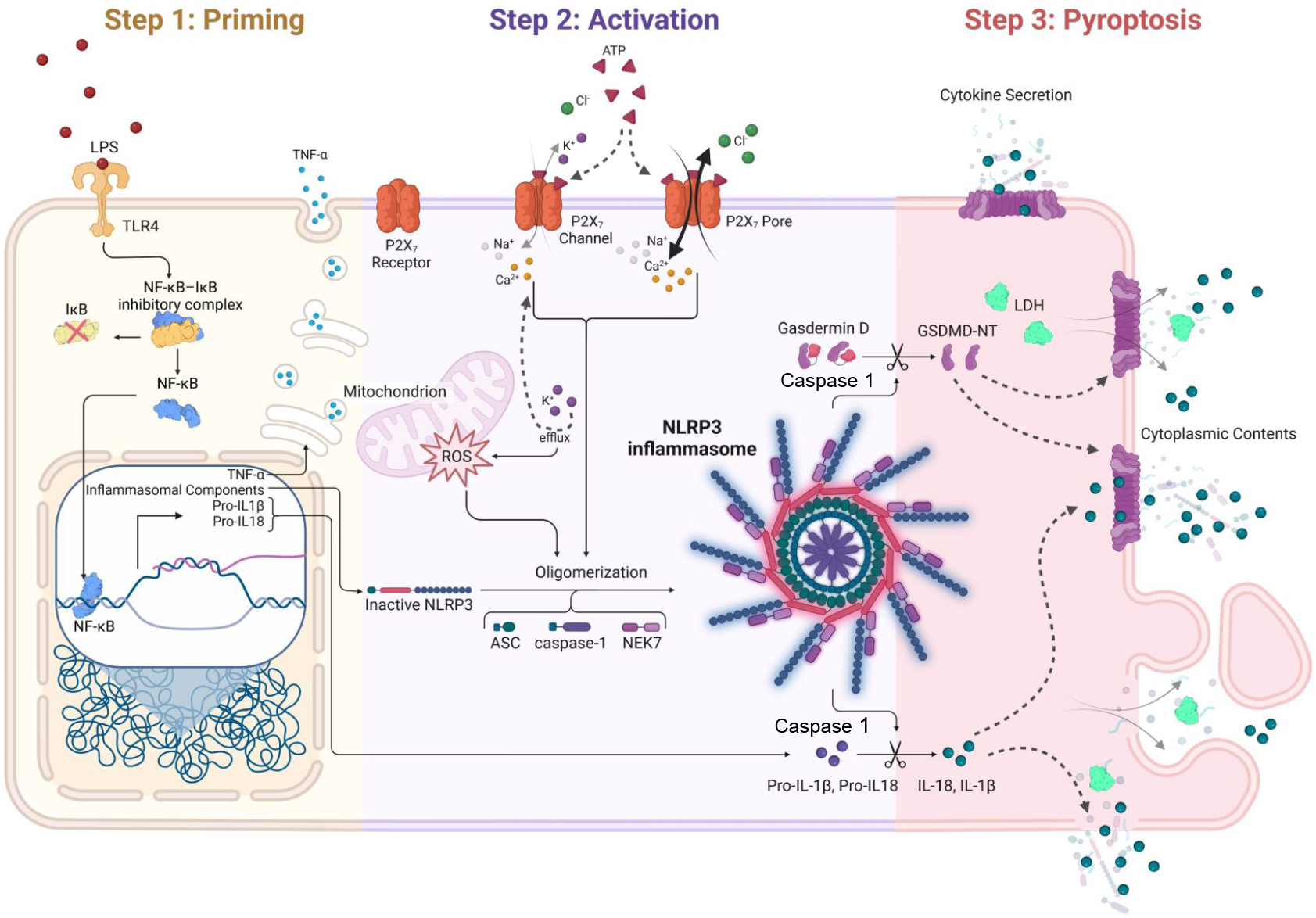
Activation of NLRP3 inflammasome. NLRP3 inflammasome needs two consecutive steps to be active: a priming stage and an activation stage. During the priming, TLR receptors can be stimulated by DAMPs, HAMPs and PAMPs which allow the translocation of nuclear factor kappa B (NF-κB) to the cell nucleus. The activation of NF-κB pathway increase the transcription of the inflammasome components, and the immature form of some pro-inflammatory cytokines. Next, a second signal such as P2X7 activation by extracellular ATP, lysosomal destabilization and/or production of ROS is required to induces the oligomerization of NLRP3 inflammasome complex which activate caspase 1. Caspase 1 can turn pro-IL-1β and pro-IL-18 into mature cytokines and cleaves the N-terminus of Gasdermin D which can oligomerizes in membranes to form pores. Liberation of pro-inflammatory cytokines such as IL-1β and IL-18, NLRP3 inflammasome complex and other cytoplasmic contents induces inflammation and a cell death type called pyroptosis.

IL-1β levels were markedly increased in the LPS and ATP-treated group compared to the control, indicating that NLRP3 activation is induced in these cells by the high-salt diet. TNF-α levels were elevated in LPS-treated groups; however, ND-13 reduced TNF-α secretion in the high-salt group, while MCC950 had no effect as we expected since inflammasome activation does not regulate NFkB activation (see **Fig. 6A** and **6B**). However, TNF-α levels were significantly decreased by effects of high salt diet, which is consistent with our results in TNF-α expression in renal cortex, supporting the hypothesis that NFkB activation is inhibited by high salt diet in our model, as have been described in previous studies (16).

**Figure 6.**
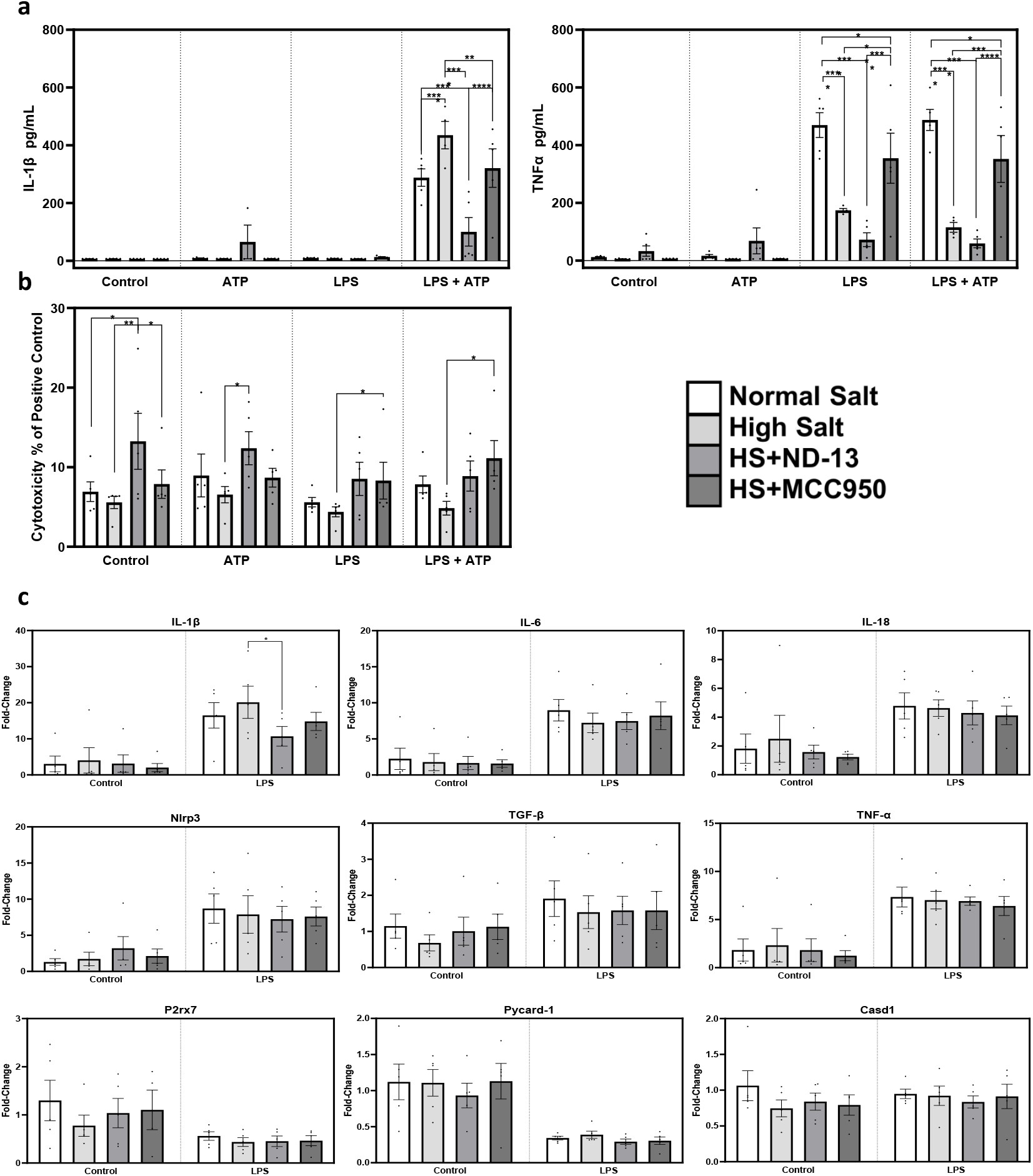
Data in peritoneal cells of mouse with high salt diet. **(a)** Release of IL-1β and TNF-α in peritoneal macrophages, (n=3-5). **(b)** Percentage of cell death compared to positive control in peritoneal macrophages, (n=3-5). **(c)** mRNA expression in peritoneal leukocytes, (n=5). Data is represented as mean ± SEM; one-way ANOVA was used in a-d; significance levels are indicated as follow: \**p<0.05*, \*\**p<0.01*, \*\*\**p<0.001*, \*\*\*\**p<0.0001*.

**Figure 7.**
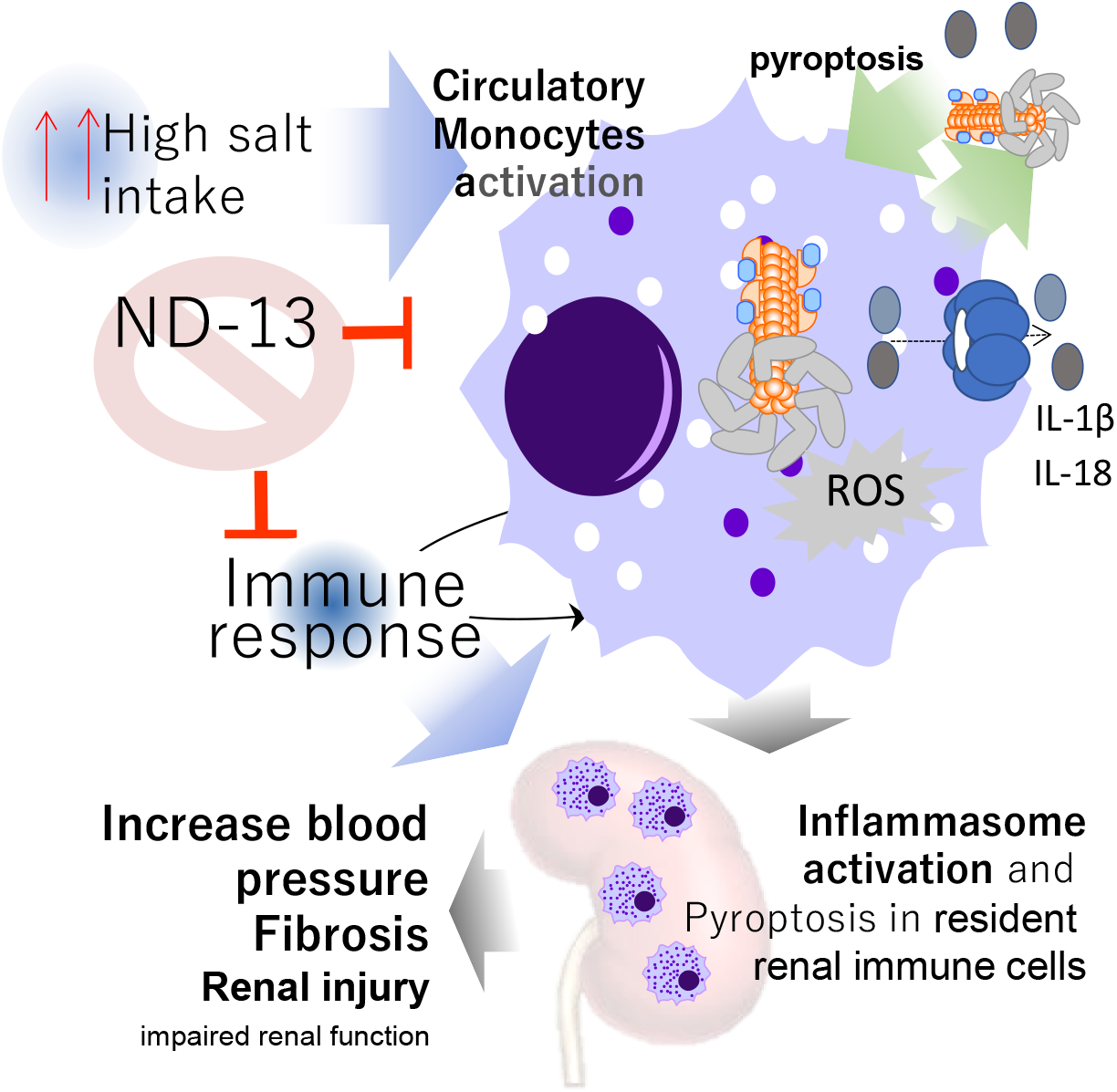
Graphical Abstract. A high-salt diet induces NLRP3 inflammasome activation in peripheral immune cells. These cells exhibit increased production of cytokines and chemokines, promoting tissue infiltration. This process contributes to renal inflammation and triggers pyroptosis in resident renal immune cells, exacerbating renal damage and elevating blood pressure.

The cytotoxicity data ruled out any potential toxic effects of the experimental treatment in the LPS/ATP group. The secreted IL-1β protein levels in each treatment group were consistent with the observed Il-1β transcript numbers. In contrast, Tnf-α mRNA exhibited much lower variability between groups compared to the observed secretion levels of its respective cytokine. Treatment with ND-13 significantly decreased Il-1β transcript levels in isolated LPS-incubated leukocytes from high-salt diet mice compared to vehicle-treated controls, in agreement with the ELISA data. However, no significant differences were observed in the expression of other interleukins and other genes associated with NLRP3 inflammasome activation indicated that gene expression itself is not a critical factor in NLRP3 activation.

These results emphasize the key role of inflammasome activation in blood pressure regulation under high-salt dietary conditions, and that gene expression in the renal cortex were not relevant on these effects (**Fig. 6**).

## 4. Discussion

This study investigates the effects of ND-13, a peptide derived from the renal DJ-1 protein, and MCC950, an NLRP3 inflammasome inhibitor, on blood pressure regulation in mice fed a high-salt diet. High salt intake led to increased blood pressure in C57Bl/6J mice, which was effectively prevented by treatments with both ND-13 and MCC950. The study found no signs of renal inflammation or collagen deposition, but proteinuria and tubular damage were observed which were prevented by ND-13 treatment. Additionally, the high-salt diet activated the NLRP3 inflammasome in peritoneal macrophages, increasing IL-1β release. Both ND-13 and MCC950 normalized inflammasome activity and reduced IL-1β expression. These findings suggest that NLRP3 inflammasome activation in peripheral immune cells plays a relevant role in blood pressure elevation, and ND-13 could represent a promising therapeutic option for preventing salt-induced hypertension and its associated inflammatory processes.

Inflammation arises from tissue damage, disruptions in homeostasis, or infection. While inflammation serves as a defense mechanism against infectious agents and supports tissue repair, it can become detrimental if prolonged (17). Decades of intensive research have demonstrated that inflammation is a key contributor to the development and progression of hypertension. For example, major inflammatory markers such as C-reactive protein (CRP), interleukin-6 (IL-6), and tumor necrosis factor-alpha (TNF-α) have been associated with an increased risk of hypertension in a meta-analysis of prospective and retrospective cohort studies (18). However, elevated blood pressure can also induce NF-κB activity, further promoting the expression of IL-6 and TNF-α (19). Current evidence, does not support the use of high-sensitivity CRP (hsCRP) to predict or prevent atherosclerotic cardiovascular diseases which are associated with development of hypertension (20). Therefore, these findings highlight the complex interplay between inflammation and blood pressure regulation. This complexity underscores the need to identify specific inflammatory markers and pathways to clarify whether inflammation is a cause or an effect of blood pressure dysregulation in clinical cases.

Inflammasomes are critical sensors of the inflammatory response. In particular, activation of the NLRP3 inflammasome in various organs, including the kidneys, vascular endothelium, and hypothalamus, contributes to the vascular and renal dysfunction observed in hypertension. For instance, in animal models of salt-induced hypertension, NLRP3 inhibition has been shown to lower blood pressure and improve renal function (21), consistent with our findings. Furthermore, recent studies have identified polymorphisms in the NLRP3 gene associated with elevated blood pressure in older populations, suggesting a genetic predisposition to hypertension mediated by inflammasome activation (5, 22).

Recent studies have evaluated the interaction between the dysregulation of epithelial sodium channels (ENaC) and NLRP3 inflammasome activation (3). Sodium dysregulation through ENaC upregulation exacerbates NLRP3 activation, increasing inflammatory cytokines such as IL-18 and IL-1β in human monocytes with cystic fibrosis-associated mutations and in human bronchial epithelial cells (23), demonstrating the effect of sodium dysregulation on NLRP3 activation. In another study, a high-sodium diet (4% NaCl) led to increased IL-1β and IL-18 production via NLRP3 activation, while inhibition of ENaC with amiloride and NLRP3 inflammasome with MCC950 mitigated these effects (24). These findings are consistent with our results and suggest a positive feedback loop between inflammasome activation and sodium retention, which triggers an increase in blood pressure and may pave the way for other cardiovascular complications (3). The role of NLRP3 in reducing blood pressure has been reported (21). However, questions remain: Is NLRP3 inflammasome overactivated under high-salt diet conditions? Is inflammasome activation the key mechanism driving blood pressure elevation, or are other pathways involved? Furthermore, previous studies did not specify the types of cells involved in NLRP3 overactivation under high-salt conditions. Notably, no changes in inflammatory markers were observed in the renal cortex in our study. Instead, only peritoneal cells from animals fed a high-salt diet showed a significant increase in NLRP3 pre-activation, which was normalized by pretreatment with ND-13 and MCC950. These findings suggest that NLRP3 inflammasome activation in circulating monocyte cells serves as the primary sensor for triggering inflammation and raising blood pressure.

ND-13, a peptide derived from the DJ-1 protein (10, 11, 25), has been shown to activate the Nrf2 pathway (26, 27), which plays a critical role in cellular defense mechanisms, particularly in response to oxidative stress and inflammation. Activation of Nrf2 upregulates antioxidant and cytoprotective genes, such as heme oxygenase-1 (HO-1) and NAD(P)H quinone dehydrogenase 1 (NQO1), which reduce oxidative damage and inflammation in tissues, including the kidneys and blood vessels (27).

In the context of hypertension, studies suggest that ND-13, through Nrf2 activation, not only protects against neurotoxic damage but also modulates inflammatory responses contributing to hypertension. For instance, ND-13 treatment has been shown to attenuate blood presse values, presumably through partial activation of Nrf2. This activation may contribute to the prevention of renal damage by mitigating inflammation (6, 8, 15, 28). Previous studies have demonstrated the potential of Nrf2 pathways to regulate blood pressure through various mechanisms, including the mitigation of oxidative stress, a key factor in hypertension (29) and the attenuation of renal inflammasome activation (4). For example, sulforaphane, an Nrf2 activator, has been shown to reduce systolic and diastolic blood pressure in hypertensive rats by enhancing antioxidant defenses (30).

Further studies are needed to elucidate the key mechanisms underlying blood pressure increases associated with high-salt conditions. In this context, NLRP3 inflammasome activation in peripheral immune cells could serve as a reliable marker for evaluating this pathology and determining appropriate treatments for each case. Given the roles of Nrf2 in blood pressure regulation and NLRP3 inflammasome activity, Nrf2 activators are being explored as potential therapeutic agents for hypertension. However, further clinical trials are essential to fully understand the efficacy and safety of Nrf2 activation in managing human hypertension.

## Acknowledgments

The authors thank the pathology and genomic platforms of the Murcia Institute of Biosanitary Research of the Region of Murcia (IMIB) for their services and the technical assistance, and IMIB-Arrixaca Biobank to store and administrate human samples. Besides, the Histology Services at the George Washington University Research Pathology Core Lab for their expertise.

## Author Contributions

SC, IA, HL, RG and PJ participated in the design of the study and in the discussion and interpretation of the data. JF addressed the animals work. RG and JP performed part of the in vitro experiments. PL to study the morphological damage. CM and MCGL performed the immunostaining data in mice. CA and EG analyzed the PAS staining data in mice. CA and SC performed the figures, the literature search and drafted the manuscript. CA figures manuscript edition for publication.

## Data sharing statement

All data, both clinical and laboratory, will be published following FAIR (Findable, Accessible, Interoperable and Reusable) principles. This will facilitate that the results of this study are easily reproducible, and that all the information generated (that does not affect sensitive data) can be easily found and reused in other studies. The human data collection will be carried out in electronic format through a Web platform (SemanticCRF) developed by the Biomedical Informatics and Bioinformatics Platform of IMIB and included in (https://dataverse.harvard.edu/).

